# Quantitive variation of male and female-specific compounds in 99 drosophilid flies

**DOI:** 10.1101/2024.06.24.600385

**Authors:** Mohammed A. Khallaf, Melissa Diaz-Morales, Bill S. Hansson, Markus Knaden

## Abstract

Variation in sex pheromones is regarded as one of the causes of reproductive isolation and speciation. We recently identified 51 male- and female-specific compounds – many of which function as sex pheromones – in 99 drosophilid species^1^. Here, we report that despite many of these compounds being shared between species, their quantities differ significantly. For example, although 34 drosophilid species share the male-specific compound cis-vaccenyl acetate (cVA), which plays a critical role in regulating various social and sexual behaviors, the amount of cVA can differ by up to 600-fold between different species. Additionally, we found 7-tricosene, the cuticular hydrocarbon pheromone, present in 35 *Drosophila* species. Our findings indicate that 7-tricosene is equally present in both sexes of 14 species, more abundant in males of 14 species, and more abundant in females of 7 species. We provide raw data on the concentration of potential pheromone components in the 99 drosophilids, which can provide important insights for further research on the behavior and evolution of these species. Quantitative variations highlight species-specific patterns, suggesting an additional mechanism for reproductive isolation built on specific combinations of compounds at set concentrations.

## Background

The diversification of sex-pheromone communication is driven by diverse factors and influenced by multiple pressures, including genetic constraints and environmental signals. Until recently, the enormous diversity of sex pheromones in *Drosophila* flies, along with their evolutionary diversification and detection, had not been comprehensibly described. We recently characterized the sex pheromone communications systems for 99 species of drosophilid flies, identifying up to 43 male-specific and 9 female-specific compounds^1^. Male-specific compounds spanned various chemical classes and were often transferred to females during mating, whereas female-specific compounds were not transferred to males. Mapping these compounds onto the phylogenetic tree showed that some male-specific compounds are widely shared across distant species, while a few are species-specific. This study highlighted how species-specific olfactory signals can reinforce sexual isolation barriers between species. However, data quantifying the abundance of these compounds for each species was previously unavailable.

## Data description

To quantify the male- and female-specific compounds, we analyzed the chemical profiles of 99 species and compared the chromatograms of both sexes within each species. Out of 99, 81 species exhibited sexually dimorphic cuticular chemicals. Remarkably, all 81 of these species showcased the presence of male-specific compounds, which amounted to a total of 42 unique compounds. In contrast, only 15 species displayed female-specific compounds, amounting to 9 compounds in total (See Table 1).

**Figure 1.**
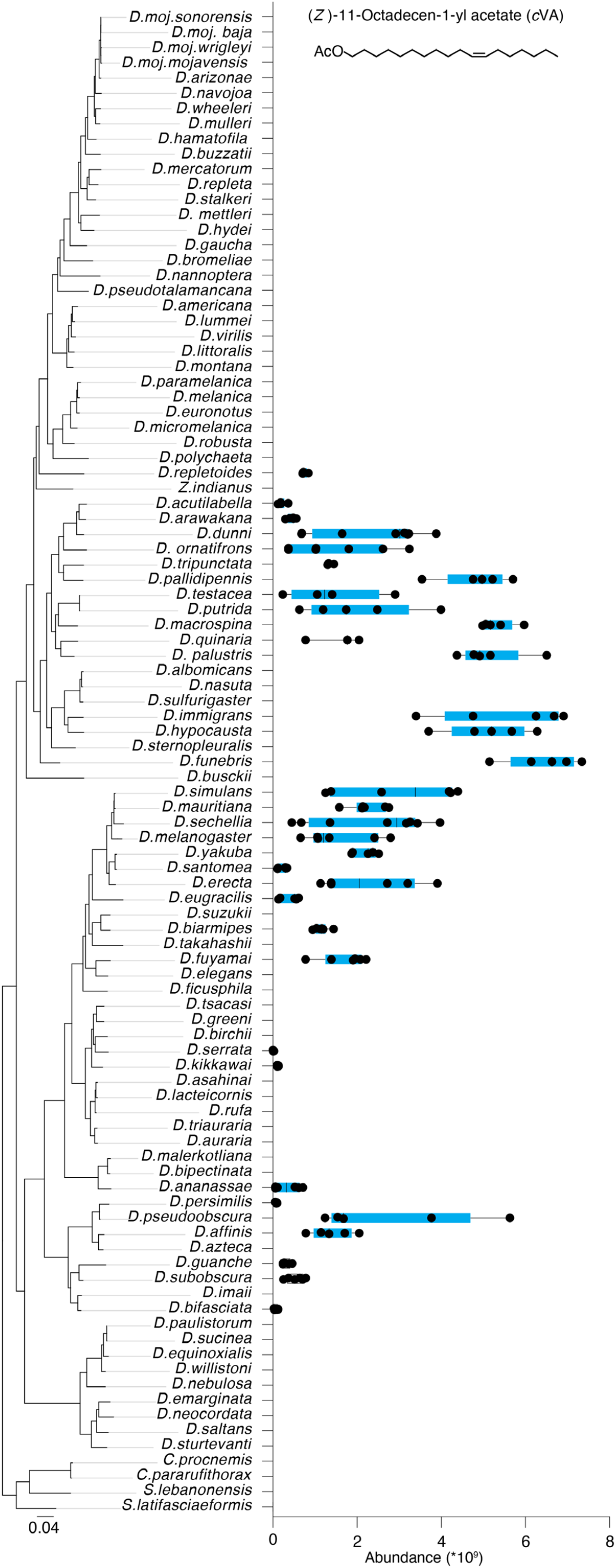
Quantification of the male-specific compound cis-vaccenyl acetate (cVA) in the 99 species. cVA was identified in 34 species, with the highest concentration observed in *D. funebris* and lowest in *D. serrata*. Each species underwent analysis with five or more replicates. Species names are ranked based on their relationships^1^. cVA is among the 42 male-specific compounds detected in the 81 dimorphic species (see Table 1).

**Figure 2.**
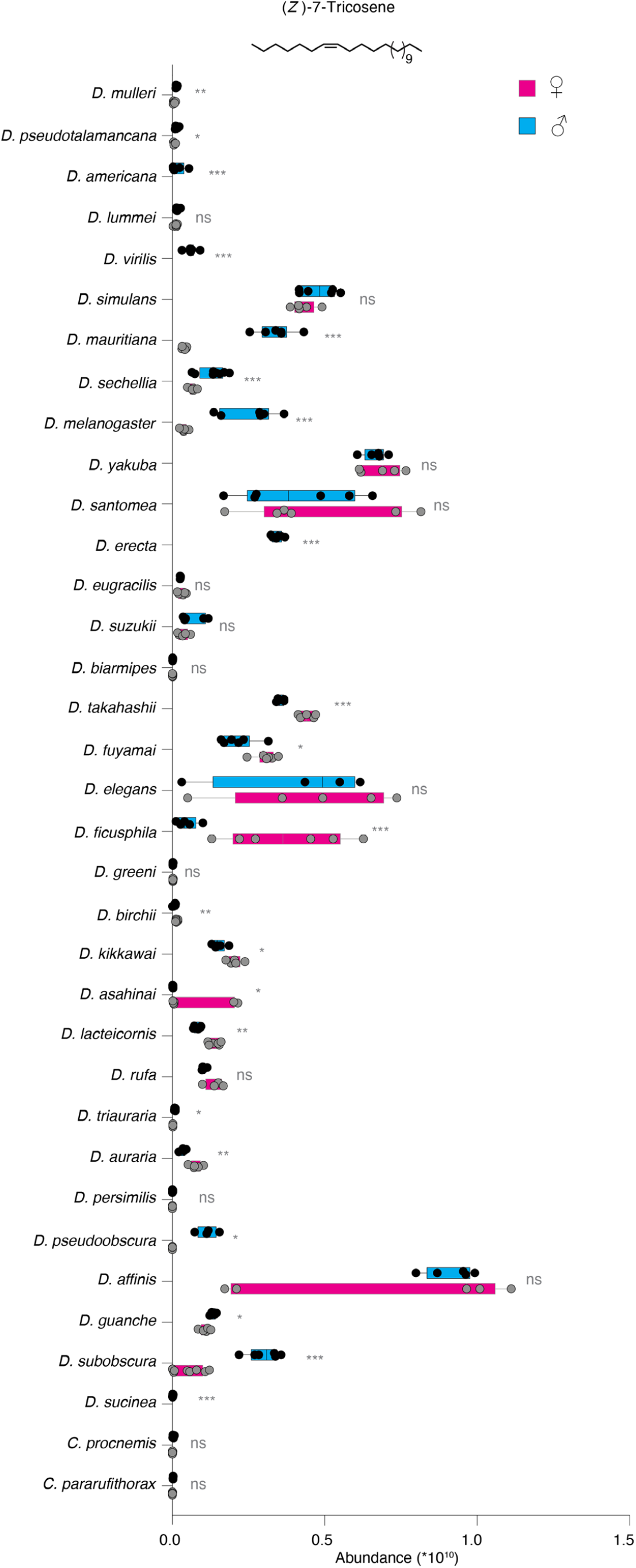
Quantification of the cuticular hydrocarbon pheromone, 7-tricosene, in 35 drosophilids. Box plots illustrate 7-tricosene abundance across five or more replicates in males (blue) and females (pink). 7-tricosene is equally present in both sexes of 14 species, more abundant in females of 7 species, and more abundant in males of 14 species. Notably, our findings indicate that 7-tricosene is a male-specific compound in four drosophilids: *D. virilis, D. americana, D. erecta* and *D. sucinea*. Pairwise comparisons between sexes within each species were conducted using the Mann-Whitney test. Ns p > 0.05; ^*^ p < 0.05; ^**^ p < 0.01; ^***^ p < 0.001.

## Experimental design, materials and methods

### Fly stocks

Wild-type flies used in this study were obtained from the National Drosophila Species Stock Centre (NDSSC; http://blogs.cornell.edu/drosophila/) and Kyoto stock center (Kyoto DGGR; https://kyotofly.kit.jp/cgi-bin/stocks/index.cgi). All flies were reared at 25 °C, 12 h Light:12 h Dark and 50% relative humidity. Stock numbers and breeding diets are listed in ^1^.

### Thermal desorption-gas chromatography-mass spectrometry (TD-GC-MS)

Individual headless vigin male and female flies in different mating status were prepared for chemical profile collection as described previously^2,3^, with some modifications. Briefly, the GC-MS device (Agilent GC 7890 A fitted with an MS 5975 C inert XL MSD unit; www.agilent.com) was equipped with an HP5-MS UI column (19091S-433UI; Agilent Technologies). After desorption at 250 °C for 3 min, the volatiles were trapped at −50 °C using liquid nitrogen for cooling. In order to transfer the components to the GC column, the vaporizer injector was heated gradually to 270 °C (12 °C/s) and held for 5 min. The temperature of the GC oven was held at 50 °C for 3 min, gradually increased (15 °C/min) to 250 °C and held for 3 min, and then to 280 °C (20 °C/min) and held for 30 min. For MS, the transfer line, source, and quad were held at 260 °C, 230 °C, and 150 °C, respectively. Eluted compounds were ionized in electron ionization (EI) source using electron beam operating at 70 eV energy and their mass spectra were recorded in positive ion mode in the range from *m/z* 33 to 500. All gas-chromatography data were collected and analyzed by MSD Chemstation software (F.01.03.2357).

### Table 1 (Supplementary file)

Quantification of the 42 male-specific compounds (Sheet 1) and 9 female-specific compounds (Sheet 2) across 99 species. Data were collected from five or more replicates of each sex, totaling over 580 samples for males and 520 for virgin females across all 99 species. Additionally, the quantification of 7-tricosene in males and females of 35 drosophilids is detailed in Sheet 3.

## CRediT author statement

**Mohammed A. Khallaf**: Conceptualization, Methodology, Investigation, Software, Data curation, Visualization, Writing-Original draft preparation, **Melissa Diaz-Morales**: Data curation, Visualization, Writing-Reviewing and Editing, **Bill Hansson**: Conceptualization, Validation, Funding acquisition, Writing-Reviewing and Editing, **Markus Knaden**: Conceptualization, Validation, Writing-Reviewing and Editing.

## Acknowledgments

We thank Ibrahim Alali for fly rearing. Wild-type flies were obtained from the San Diego Drosophila Species Stock Center (now The National Drosophila Species Stock Center, Cornell University) and KYOTO Stock Center. This research was supported through funding by the Max Planck Society.

